# HNF4α-TET2-FBP1 axis contributes to gluconeogenesis and type 2 diabetes

**DOI:** 10.1101/2024.09.29.615677

**Authors:** Hongchen Li, Xinchao Zhang, Xiaoben Liang, Shuyan Li, Ziyi Cui, Xinyu Zhao, Kai Wang, Bingbing Zha, Haijie Ma, Ming Xu, Lei Lv, Yanping Xu

## Abstract

The control of gluconeogenesis is critical for glucose homeostasis and the pathology of type 2 diabetes (T2D). Here, we uncover a novel function of TET2 in the regulation of gluconeogenesis. In mice, both fasting and a high-fat diet (HFD) stimulate the expression of TET2, and TET2 knockout impairs glucose production. Mechanistically, FBP1, a rate-limiting enzyme in gluconeogenesis, is positively regulated by TET2 in liver cells. TET2 is recruited by HNF4α, contributing to the demethylation of FBP1 promoter and activating its expression in response to glucagon stimulation. Moreover, metformin treatment increases the phosphorylation of HNF4α on Ser313, which prevents its interaction with TET2, thereby decreasing the expression level of FBP1 and ameliorating the pathology of T2D. Collectively, we identify an HNF4α-TET2-FBP1 axis in the control of gluconeogenesis, which contributes to the therapeutic effect of metformin on T2D and provides a potential target for the clinical treatment of T2D.

## Introduction

Loss of glucose homeostasis can lead to type 2 diabetes (T2D), characterized by persistently elevated blood glucose levels that may result in various complications, including kidney failure and neuropathy. Since abnormal gluconeogenic activity is the primary contributor to hepatic glucose production (HGP) (1, 2), which is the main source of increased blood glucose concentration in T2D, targeting gluconeogenesis represents a viable strategy for maintaining blood glucose homeostasis.

Fructose 1,6-bisphosphatase (FBP1), a rate-controlling enzyme in gluconeogenesis that catalyzes the conversion of fructose 1,6-bisphosphate to fructose 6-phosphate. Notably, a point mutation in FBP1 has been reported to significantly reduce the efficacy of metformin, a first-line drug for the treatment of T2D, suggesting that FBP1 plays a major role in the therapeutic effect of metformin (3). However, the specific response of FBP1 to metformin treatment remains unclear.

Ten-Eleven Translocation 2 (TET2) belongs to the TET family, which includes TET1, TET2, and TET3. These enzymes function as DNA dioxygenases, catalyzing the successive oxidation of 5-methylcytosine (5mC) to 5-hydroxymethylcytosine (5hmC) (4), 5-formylcytosine (5fC), and 5-carboxycytosine (5caC) (5, 6). Subsequently, thymine-DNA glycosylase (TDG) mediates the removal of 5caC, resulting in the formation of an unmodified cytosine, which participates in DNA demethylation and gene expression regulation (6). Recently, it was reported that the expression of FBP1 can be regulated by promoter methylation (7, 8), leading us to hypothesize that TET2 may play a role in regulating FBP1 expression and gluconeogenesis.

Mutations in TET2 frequently occur in various myeloid cancers. Somatic alterations in TET2 are observed in 50% of patients with chronic myelomonocytic leukemia and are associated with poor outcomes (9). The frequency of TET2 mutations in patients with myelodysplastic syndromes is 19% (10). Moreover, reversing TET2 deficiency suppresses the abnormal differentiation and self-renewal of hematopoietic stem and progenitor cells (HSPCs), and blocks leukemia progression (11). Additionally, TET2 has also been reported to repress mTORC1 and HIF signaling, thereby suppressing tumor growth in hepatocellular carcinoma (HCC) and clear cell renal cell carcinoma (ccRCC), respectively (12, 13). These studies focused on the function of TET2 in cancer development, however, it is unclear whether TET2 is involved in T2D progression. In this study, we demonstrated that TET2 is recruited by HNF4α to the FBP1 promoter, activating FBP1 expression through demethylation, which contributes to gluconeogenesis and T2D pathology. Furthermore, we identified HNF4α-TET2-FBP1 axis as a target of metformin treatment, suggesting that targeting this axis may represent a potential strategy for T2D management.

## Results

### TET2 contributes to gluconeogenesis and T2D

To investigate the role of TET2 in gluconeogenesis, we developed three mouse models: one subjected to overnight fasting for 16 hours prior to testing and two subjected to high-fat feeding (HFD) for 11 days and 12 weeks, respectively. The results indicated that both fasting and HFD increased the mRNA and protein levels of Tet2 in the livers of mice compared to the normal chow group (Fig. 1A-E), suggesting that TET2 may play an important role in gluconeogenesis. To test this hypothesis, we examined the effect of TET2 on glucose output and found that TET2 overexpression promoted glucose output in HepG2 cells and primary mouse hepatocytes (Fig. 2A and B). Consistent with this, *TET2* knockout impaired gluconeogenesis in HepG2 cells and mouse hepatocytes, even under glucagon treatment (Fig. 2C and D). Collectively, these data demonstrated that TET2 contributes to gluconeogenesis, prompting us to investigate whether TET2 is involved in T2D progression. Pyruvate tolerance test (PTT), glucose tolerance test (GTT), and insulin tolerance test (ITT) were conducted, revealing *Tet2* KO significantly increased glucose tolerance and insulin sensitivity compared to the control mice (Fig. 2E-H). Additionally, to assess the plasma insulin levels in response to GTT in *Tet2-*KO mice, we collected the orbital venous blood and examined the insulin levels at different time point. The results showed that *Tet2-KO* mice exhibited lower insulin secretion after glucose administration, further reflecting higher insulin sensitivity (Fig. 2H). Meanwhile, we compared the body weight differences between wild type (WT) and *Tet2*-KO mice and found no significant differences at 8 and 10 weeks of age on normal chow (Fig. 2I). In summary, the loss of *Tet2* function decreased gluconeogenesis in the liver and may contribute to the treatment of T2D.

**Figure 1.**
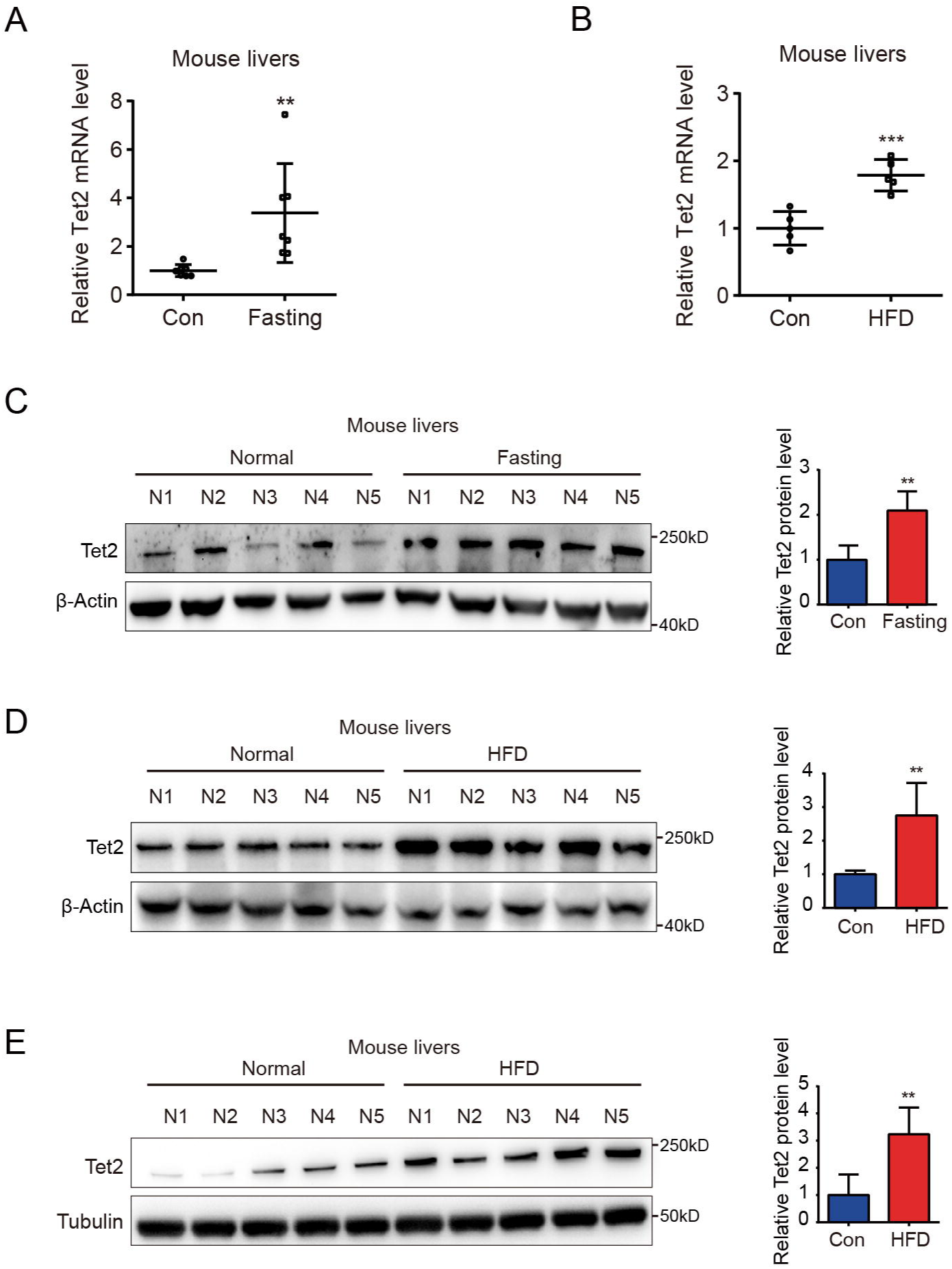
TET2 expression increases in fasting and HFD mouse livers. (**A**) qRT-PCR analysis of Tet2 mRNA levels in mouse livers after 16 h fasting treatment. The data are normalized to the GAPDH expression (n =7). (**B**) qRT-PCR analysis of Tet2 mRNA levels in mouse livers after high-fat diet treatment for 11 days. The data are normalized to the GAPDH expression (n=7). (**C**) Western blot analysis and quantification of Tet2 protein levels in mouse livers following 16 h fasting treatment (n=5). (**D**) Western blot analysis and quantification of Tet2 protein levels in mouse livers following the high-fat diet treatment for 11 days (n=5). (**E**) Western blot analysis and quantification of Tet2 protein levels in mouse livers following the 12-week high-fat diet treatment (n=5).

**Figure 2.**
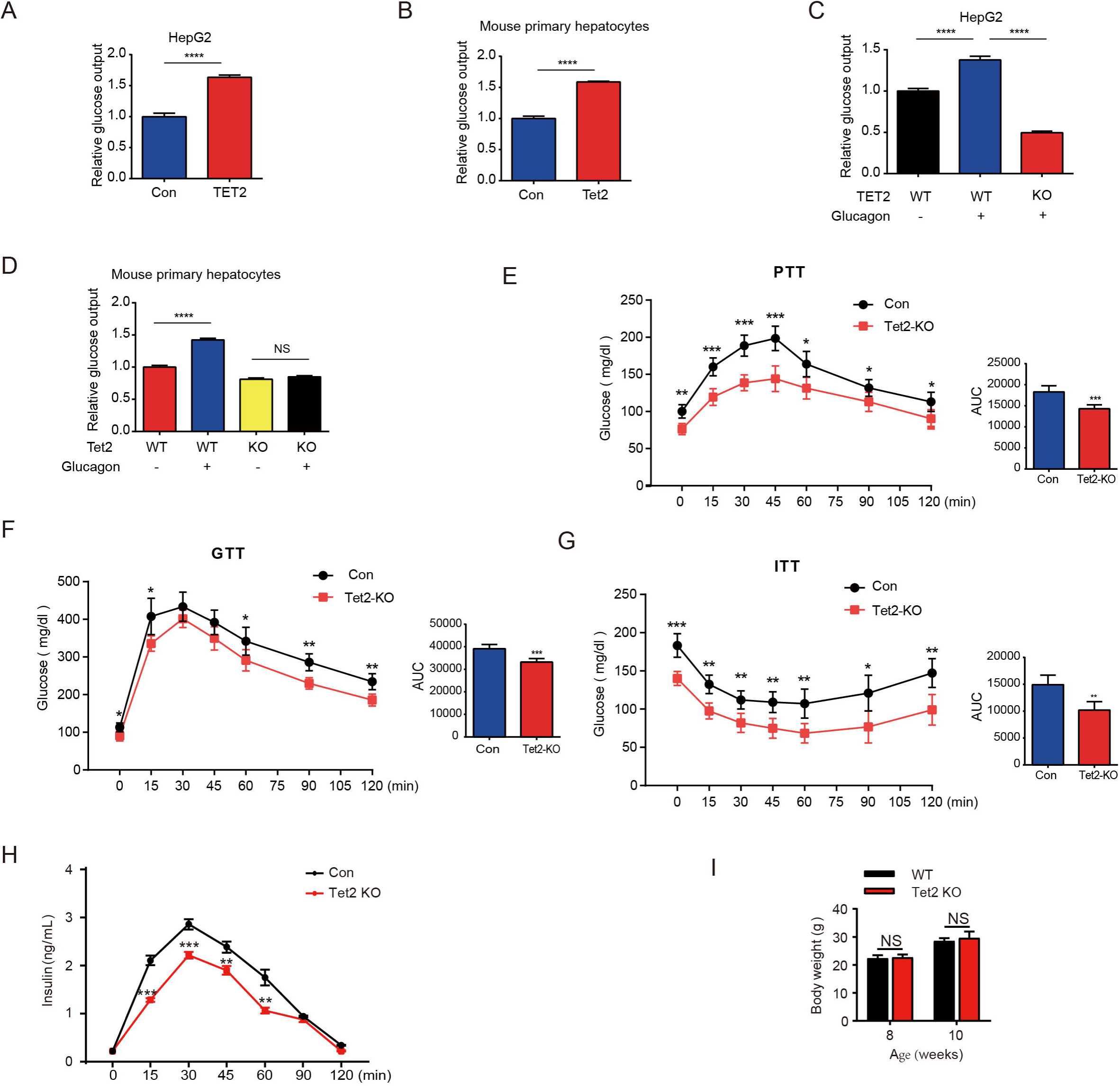
TET2 boosts gluconeogenesis. (**A**) Glucose production assays were performed in HepG2 cells after TET2 overexpression. Data are represented as the mean±SD (n=3). (**B**) Glucose production assays were performed in mouse primary hepatocytes after Tet2 overexpression. Data are represented as the mean ± SD (n=3). (**C**) Glucose production assays were performed in HepG2 cells pre-treated with 20nM glucagon in WT and TET2 KO HepG2 cells. Data are represented as the mean ± SD (n=3). (**D**) Glucose production assays were performed in mouse primary hepatocytes pre-treated with 20nM glucagon in WT and *Tet2* KO cells. Data are represented as the mean ± SD (n=3). (**E**) PTT was performed following a 16 h fasting treatment and intraperitoneal (i.p.) injection of 1 g/kg sodium pyruvate (n = 5). (**F**) GTT was performed after a 12 h fasting treatment and i.p. injection of 2 g/kg glucose (n = 5). (**G**) ITT was performed after a 4 h fasting treatment and i.p. injection of 0.75 U/kg insulin (n = 5). (**H**) Glucose-stimulated insulin secretion was examined. After fasting and i.p. injection of 2 g/kg glucose, plasma insulin levels were measured at the indicated time points (n = 5). **(I)** Body weight of 8 or 10-week-old male WT mice and *Tet2* KO mice on a normal chow diet (n = 5).

### TET2 upregulates FBP1 expression in liver cells

Next, we explored the potential mechanism by which TET2 increases gluconeogenesis. Fructose-1,6-bisphosphatase 1 (FBP1), a rate-limiting enzyme in gluconeogenesis, plays a crucial role in T2D and was recently identified as a target of metformin (3). This led us to hypothesize that FBP1 might participate in TET2-mediated regulation of gluconeogenesis. The results showed that glucagon significantly increased both TET2 and FBP1 expression levels in HepG2 and primary mouse liver cells (Fig. 3A-C), while had no effect on the expression of PEPCK and G6Pase, other two key gluconeogenesis genes (Fig. 3D), suggesting that TET2 may regulate gluconeogenesis via FBP1. To assess the long-term effects of a single dose glucagon, we examined TET2 and FBP1 mRNA levels in different times in HepG2 cells. Interestingly, the results showed that the expression peak of TET2 and FBP1 mRNA levels occurred 30 minutes after glucagon treatment, with the prolonged effects of glucagon on TET2 mRNA lasting for more than 48 hours (Fig. 3E), indicating TET2-FBP1 axis may play an important role in gluconeogenesis in response to glucagon. Furthermore, fasting and high-fat diet (HFD) also upregulated Fbp1 mRNA levels in mouse livers (Fig. 3F and G). Notably, Pearson correlation analysis revealed a positive correlation between Tet2 and Fbp1 expression in both control and fasting groups (Fig. 3H). To confirm this, Gene Expression Profiling Interactive Analysis (GEPIA) (14) was used to analyze the correlation between TET2 and FBP1 expression in human liver tissue. Consistent with the mouse data, FBP1 expression levels positively correlated with TET2 levels in human livers (Fig. 3I). These findings prompted us to examine whether TET2 regulates FBP1 expression. The results showed that TET2 overexpression promoted FBP1 expression in primary mouse hepatocytes and HepG2 cells (Fig. 3J), while *TET2* KO significantly decreased FBP1 levels in HepG2 and LO-2 cells (Fig. 3K and L). Moreover, *TET2* KO abolished the glucagon-induced upregulation of FBP1 (Fig. 3M), suggesting that TET2 is required for glucagon-induced FBP1 upregulation. Taken together, these data indicate that TET2 regulates FBP1 expression in liver cells.

**Figure 3.**
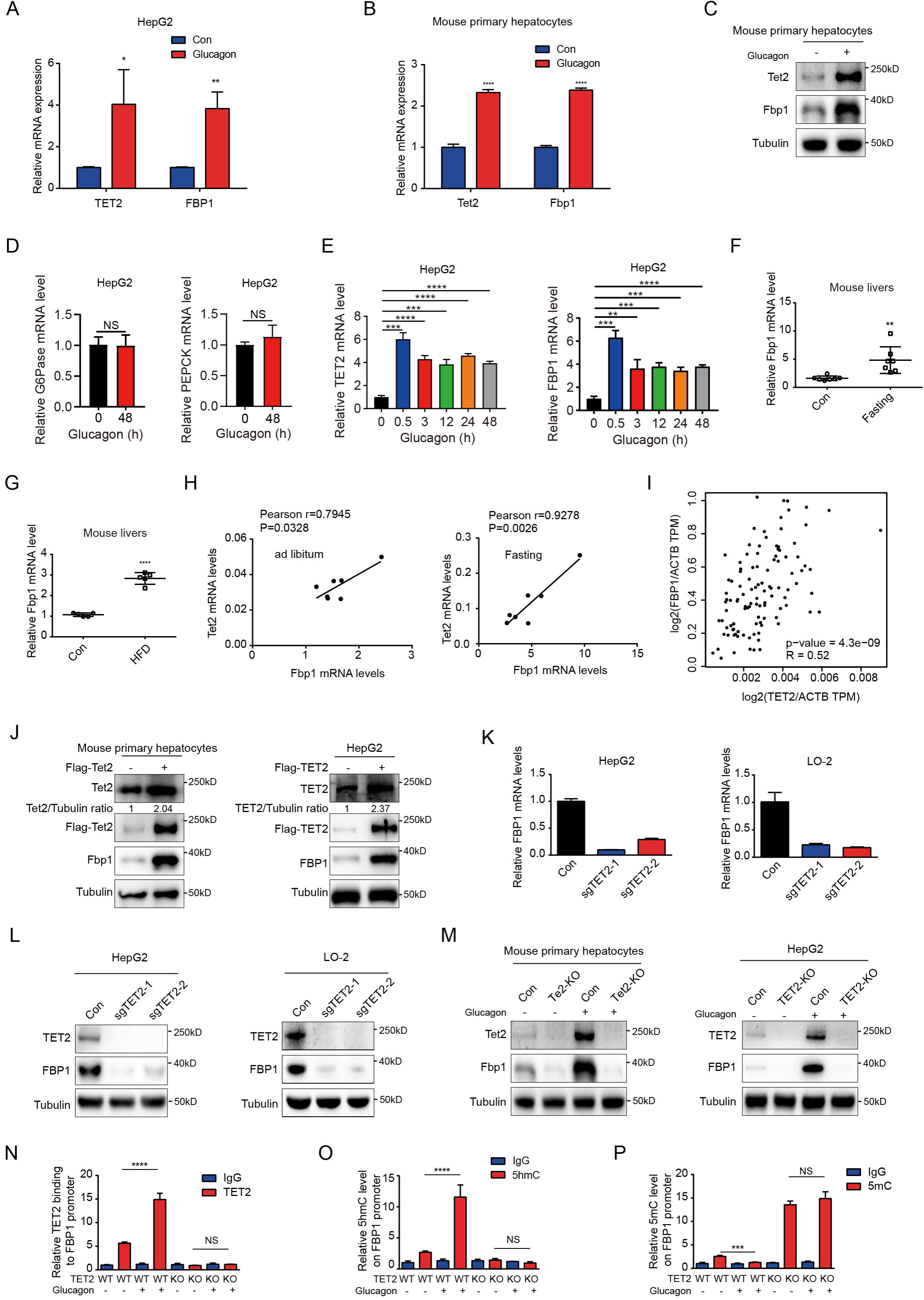
TET2 up-regulates FBP1 expression in liver cells. (**A**) qPCR analysis of TET2 and FBP1 mRNA expression levels after 20nM glucagon treatment for 48 h in HepG2 cells. Data are represented as the mean ± SD (n=3). (**B**) qPCR analysis of Tet2 and Fbp1 mRNA expression levels after 20nM glucagon treatment for 48 hours in primary mouse hepatocytes. Data are represented as the mean ± SD (n=3). (**C**) Western blot analysis of Tet2 and Fbp1 protein levels after 20nM glucagon treatment in mouse primary hepatocyte cells. (**D**) qPCR analysis of G6Pase and PEPCK mRNA expression levels after 20nM glucagon treatment for 48 hours in HepG2 cells. Data are represented as the mean ± SD (n=3). (**E**) qPCR analysis of TET2 and FBP1 mRNA expression levels after 20nM glucagon treatment at the indicated time points (n = 5). (**F**) qPCR analysis of Fbp1 mRNA levels in mouse livers following fasting treatment (n=7). (**G**) qPCR analysis of Fbp1 mRNA levels in mouse livers following HFD treatment (n=5). (**H**) Correlation analysis between Tet2 and Fbp1 levels using data from Figure 1A in mouse livers with or without fasting treatment. (**I**) Correlation analysis between TET2 and FBP1 levels in human livers. Data were collected from GEPIA(29). (**J**) Western blot analysis of TET2 and FBP1 expression after overexpression of Flag-TET2 in mouse primary hepatocytes and HepG2 cells. (**K**) qPCR analysis of FBP1 expression levels in control and TET2 knockout HepG2 and LO-2 cells. (**L**) Western blot analysis of TET2 and FBP1 protein levels in control and TET2 knockout HepG2 and LO-2 cells. (**M**) Western blot analysis of Tet2 and Fbp1 protein levels in control and Tet2 knockout mouse primary hepatocytes and HepG2 cells treated with or without 20nM glucagon. (**N**) ChIP-qPCR analysis of TET2 binding to FBP1 promoter in response to glucagon stimulation in control and TET2 knockout HepG2 cells. (**O**) ChIP-qPCR analysis of 5hmC levels in FBP1 promoter in response to glucagon stimulation in control and TET2 knockout HepG2 cells. (**P**) ChIP-qPCR analysis of 5mC levels in FBP1 promoter in response to glucagon stimulation in control and TET2 knockout HepG2 cells.

To explore the mechanism by which TET2 regulates FBP1, we performed ChIP-qPCR to investigate whether TET2 binds to the FBP1 promoter and catalyzes the conversion of 5mC to 5hmC in HepG2 cells under glucagon treatment. We found that glucagon treatment promoted TET2 binding to the FBP1 promoter in HepG2 cells, increasing 5hmC levels and reducing 5mC levels (Fig. 3N-P). Importantly, *TET2* knockout blocked this process and led to the accumulation of 5mC in FBP1 promoter (Fig. 3N-P). These data demonstrate that TET2 mediates the transcriptional activation of FBP1 in response to glucagon stimulation.

### HNF4α is required for TET2 mediated transcriptional activation of FBP1

Given that TET2 binds to DNA without sequence specificity, we sought to understand how TET2 specifically activates FBP1 expression in response to glucagon treatment. Recent genome-wide methylation and transcriptome analyses identified hepatocyte nuclear factor 4 alpha (HNF4α) as a master gluconeogenic transcription factor that plays a critical role in the pathogenesis of diabetic hyperglycemia (15). Importantly, ChIP-seq data suggest that HNF4α binds to the FBP1 promoter and participates in the regulation of gluconeogenesis in adult mouse hepatocytes (16, 17). To determine whether HNF4α is involved in TET2-mediated FBP1 expression, we performed immunofluorescence to examine the colocalization of TET2 and HNF4α with or without glucagon treatment in HepG2 cells. The results indicated that the co-localization of TET2 and HNF4α significantly increased upon glucagon treatment (Fig. 4A). Moreover, fasting and HFD treatments also promoted the co-localization of Tet2 and Hnf4α in mouse hepatocyte cells (Fig. 4B and C). Consistently, we observed that glucagon increased the interaction between TET2 and HNF4α in HepG2 cells (Fig. 4D). To determine the function of HNF4α in TET2 mediated regulation of FBP1 expression, we knocked down HNF4α in HepG2 cells using siRNA (Fig. 4E), and found that HNF4α knockdown significantly inhibited glucagon-induced TET2 binding to the FBP1 promoter (Fig. 4F), decreased 5hmC levels (Fig. 4G) and increased 5mC levels in FBP1 promoter (Fig. 4H), thereby suppressing TET2-mediated FBP1 expression (Fig. 4I) and impairing glucose output under glucagon treatment (Fig. 4J), demonstrating that HNF4α recruits TET2 to the FBP1 promoter and activates FBP1 expression through demethylation to facilitate gluconeogenesis. Notably, the expression of both Hnf4α and Fbp1 also increased in mouse livers under fasting or HFD treatment compared to the control group (Fig. 4K-M). In conclusion, these results support the notion that TET2-mediated FBP1 expression and gluconeogenesis is dependent on HNF4α.

**Figure 4.**
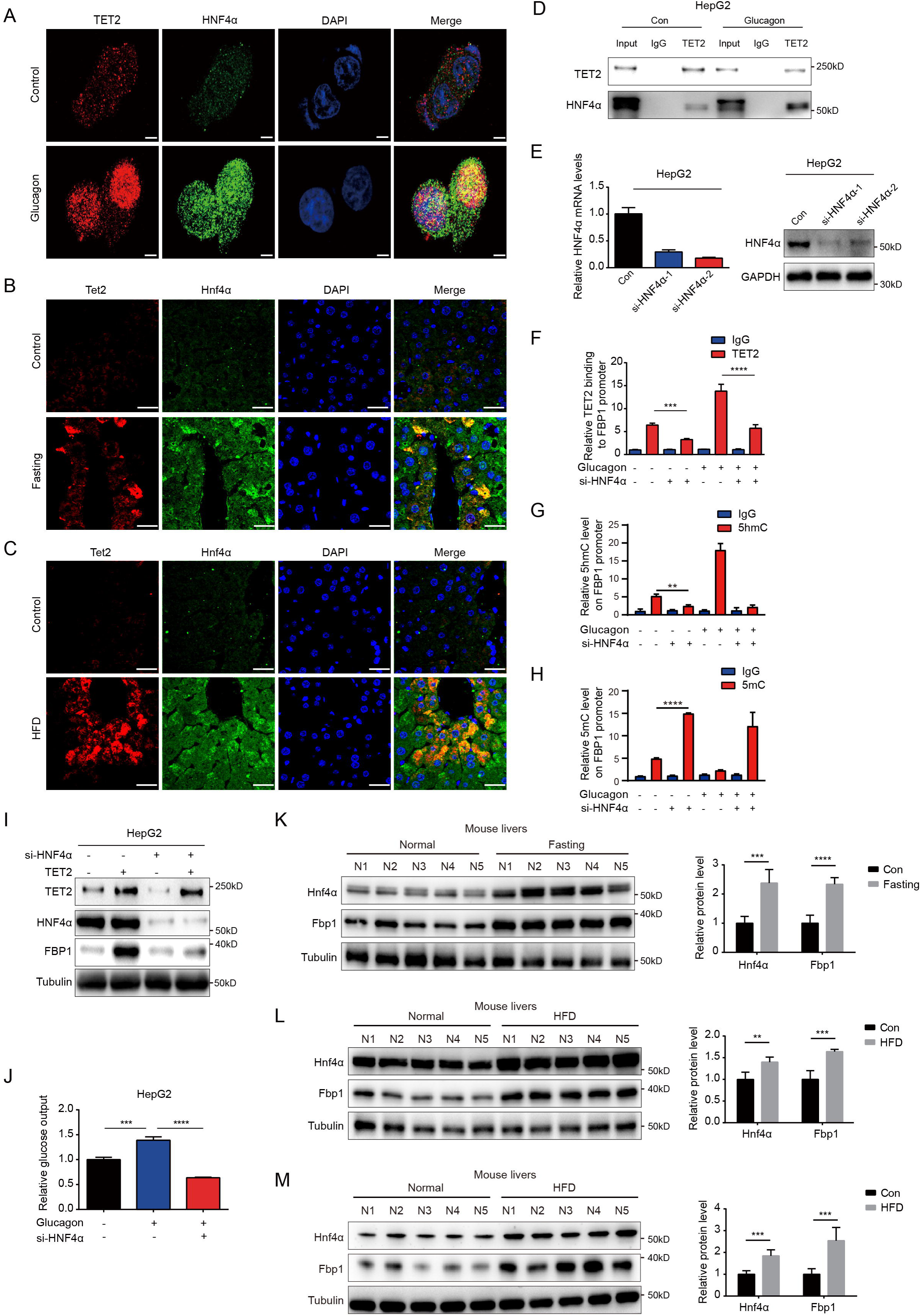
HNF4α is necessary for TET2 mediated FBP1 up-regulation. (**A**) Immunofluorescence analysis of TET2 and HNF4α co-localization in HepG2 cells after 20nM glucagon treatment for 48 h. Scale bar: 10μm. (**B**) Immunofluorescence analysis of Tet2 and Hnf4α co-localization in liver sections from standard chow and fasting mice. Scale bar: 30μm. (**C**) Immunofluorescence analysis of Tet2 and Hnf4α co-localization in liver sections from standard chow and HFD mice. Scale bar: 30μm. (**D**) Endogenous co-immunoprecipitation followed by western blot analysis of the interaction between HNF4α and TET2 with or without glucagon treatment in HepG2 cells. (**E**)qRT-PCR and western blot analysis of HNF4α expression levels in HepG2 cells transfected with two specific siRNAs. (**F**) ChIP-qPCR analysis of TET2 binding to FBP1 promoter in HepG2 cells treated with siRNA targeting HNF4α and glucagon as indicated. (**G**) ChIP-qPCR analysis of 5hmC levels in FBP1 promoter in HepG2 cells treated with siRNA targeting HNF4α and glucagon as indicated. (**H**) ChIP-qPCR analysis of 5mC levels in FBP1 promoter in HepG2 cells treated with siRNA targeting HNF4α and glucagon as indicated. (**I**) Western blot analysis of FBP1 protein levels in HepG2 cells treated with TET2 overexpression and siRNA targeting HNF4α as indicated. (**J**) Glucose production assays were performed in HepG2 cells treated with glucagon and transfected with HNF4α siRNA as indicated. (**K**-**M)**, Western blot analysis and quantification of Hnf4α and Fbp1 protein levels in mouse livers from the mice treated with 16 h overnight fasting (K) or 11-day HFD (L), or 12-week HFD treatment (M). n=5.

### HNF4α phosphorylation affects its binding to TET2 and FBP1 expression

Our results demonstrated that HNF4α recruits TET2 to the FBP1 promoter and activates FBP1 expression through demethylation, playing a crucial role in the regulation of hepatic glucose output. However, it remains unclear whether the HNF4α-TET2-FBP1 axis responds to metformin treatment. Metformin, a first-line antidiabetic drug widely used to treat hyperglycemia in type 2 diabetes, is also a well-established adenosine 5’-monophosphate-activated protein kinase (AMPK) activator (18). Interestingly, one study revealed that AMPK phosphorylates HNF4α at Ser 313, reducing its transcriptional activity (19). Combining these studies with our findings, we wonder whether metformin can affect the HNF4α-TET2-FBP1 axis. To explore the role of metformin-induced AMPK phosphorylation of HNF4α in FBP1 expression, we treated cells with metformin and assessed the interaction between HNF4α and TET2. The results showed that metformin administration impaired HNF4α’s ability to bind to TET2 (Fig. 5A), leading to a significant reduction in TET2 binding to the FBP1 promoter (Fig. 5B). Consistently, metformin treatment significantly decreased the expression level of FBP1 (Fig. 5C). Notably, metformin also induced high levels of HNF4α phosphorylation at Ser 313 (Fig. 5C). To determine the effect of HNF4α phosphorylation on HNF4α-TET2-FBP1 axis, we transfected HepG2 cells with wild type HNF4α and Ser 313 mutants. The results showed that the phosphomimetic mutation (S313D) of HNF4α impaired its ability to bind to TET2 (Fig. 5D), prevented TET2 from binding to the FBP1 promoter (Fig. 5E), and reduced FBP1 expression at both mRNA and protein levels (Fig. 5F). In contrast, the phosphoresistant mutation (S313A) showed higher activity in interacting with TET2, recruiting TET2 to the FBP1 promoter, and activating its expression (Fig. 5D-F). Taken together, these data demonstrate that metformin-mediated HNF4α phosphorylation suppresses FBP1 expression by preventing TET2 recruitment to the FBP1 promoter by HNF4α.

**Figure 5.**
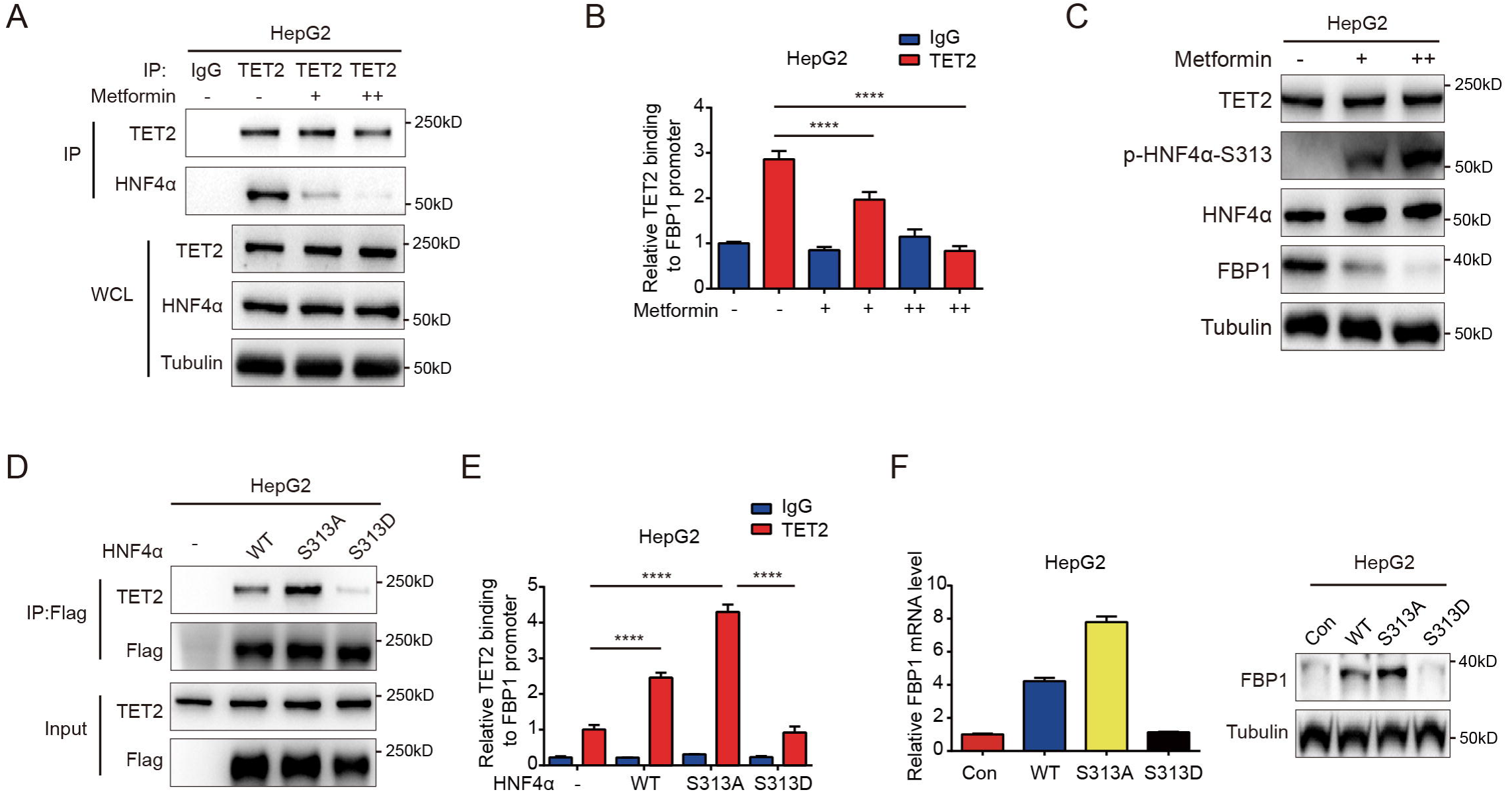
Metformin impairs the ability of HNF4α binding to TET2 and FBP1 expression. (**A**) Endogenous co-immunoprecipitation followed by western blot analysis of the interaction between HNF4α and TET2 with or without metformin (10 mM) treatment in HepG2 cells. (**B**) ChIP-qPCR analysis of TET2 binding to FBP1 promoter in HepG2 cells treated with or without metformin (10 mM). (**C**) Western blot analysis of TET2, HNF4α, HNF4α phosphorylation at Ser 313 and FBP1 levels in HepG2 cells treated with metformin as indicated (+: 5 mM, ++: 10 mM). (**D**) Endogenous co-immunoprecipitation followed by western blot analysis of the interaction between TET2 and HNF4α wildtype and S313 mutants as indicated. (**E**) ChIP-qPCR analysis of TET2 binding to FBP1 promoter in HepG2 cells transfected with HNF4α wildtype and S313 mutants as indicated. (**F**) qRT-PCR and western blot analysis of FBP1 levels in HepG2 cells transfected with HNF4α wildtype and S313 mutants as indicated.

### Targeting TET2 improves the efficacy of metformin in glucose metabolism *in vivo*

Increased rates of gluconeogenesis, derived from continuously excessive hepatic glucose production (HGP), result in abnormal glucose homeostasis T2D. Fasting or high-fat diet (HFD)-induced T2D mice showed elevated levels of Tet2 and Fbp1 in hepatocytes (Fig. 1A-E). However, it remains unclear whether knockdown of *Tet2*, in combination with metformin, would decrease glucose production, thereby improving T2D treatment outcomes. HFD-induced diabetic mice were infused with AAV8 or AAV8-sh*Tet2*. To confirm the efficiency of liver-specific *Tet2* knockdown *in vivo*, we examined *Tet2* mRNA levels in mouse livers, and found that TET2 expression significantly decreased upon AAV8-sh*Tet2* treatment (Fig. 6A). Notably, fasting blood glucose and insulin levels were markedly lower in AAV8-sh*Tet2* group than the control group in response to metformin treatment (Fig. 6B and C). PTT performed on HFD mice indicated that Tet2 suppression sharply decreased hepatic gluconeogenesis (Fig. 6D), which was consistent with lower protein levels of Fbp1 in mouse livers (Fig. 6E). Moreover, GTT and ITT showed that *Tet2* knockdown in HFD mice significantly enhanced glucose tolerance and insulin sensitivity compared to control group (Fig. 6F and G). Of note, the beneficial effects of *Tet2* knockdown alone were comparable to those of metformin in lowering glucose levels and improving insulin sensitivity. Interestingly, the combination of metformin and *Tet2* knockdown in HFD mice exhibited better glucose-lowering effects and insulin sensitivity compared to either *Tet2* knockdown or metformin. Collectively, these data demonstrated that suppressing TET2 synergizes with metformin to lower glucose production and enhance insulin sensitivity by inhibiting FBP1 expression.

**Figure 6.**
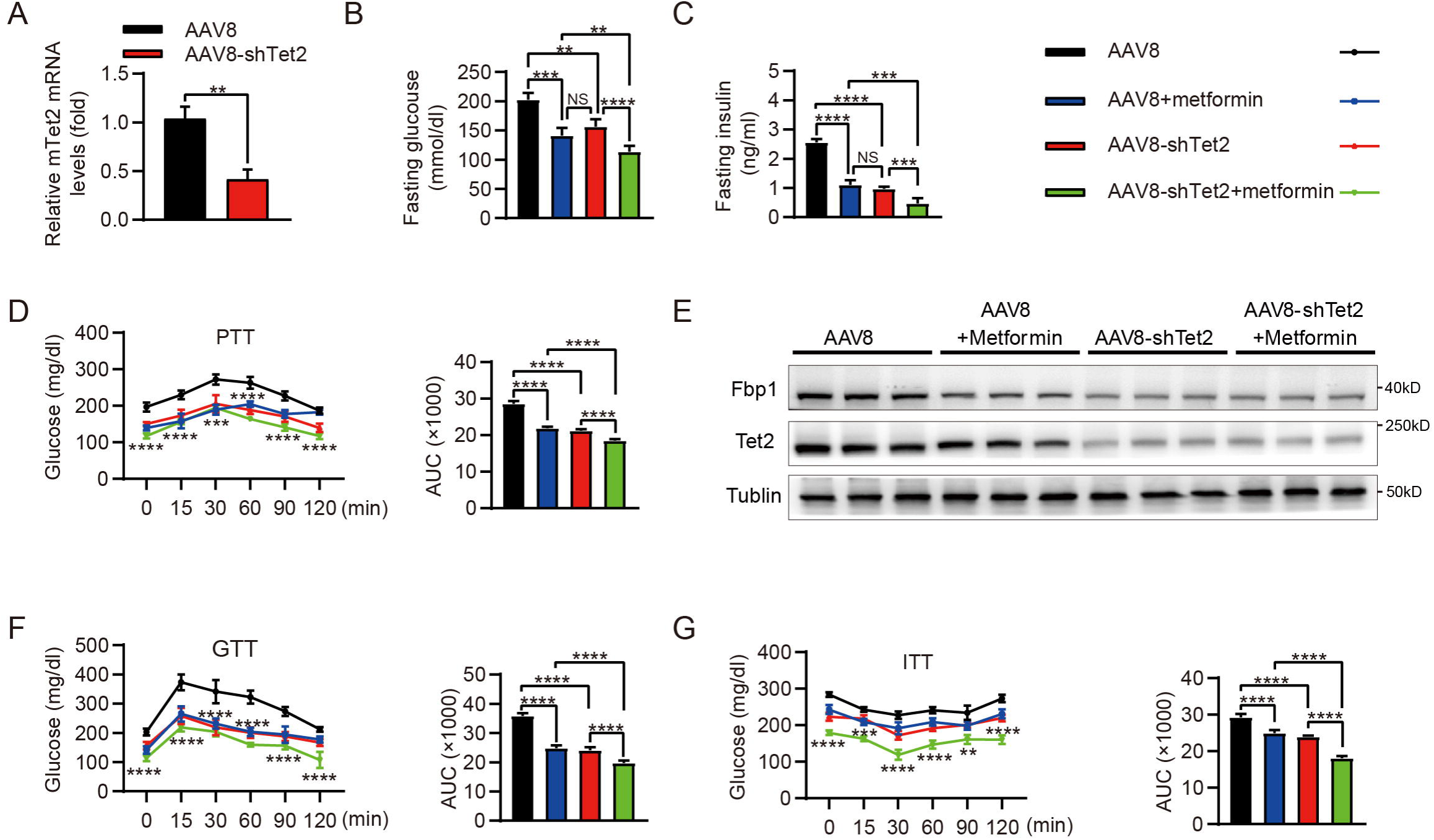
Targeting TET2 improves the efficacy of metformin in glucose metabolism *in vivo*. (**A**) qRT-PCR analysis of *Tet2* mRNA levels in mouse livers from the HFD mice infected with AAV8 or AAV8-*shTet2* for 10 days. (**B**, **C**) Analysis of fasting blood glucose (B) and plasma insulin (C) levels in the HFD mice infected with AAV8 or AAV8-*shTet2* for 10 days, and treated with or without metformin (300 mg/kg/day) for another 10 days as indicated. n = 6. (**D)** PTT was performed in the HFD mice infected with AAV8 or AAV8-*shTet2* for 10 days, and treated with or without metformin (300 mg/kg/day) for another 10 days as indicated (n = 6). (**E)** Western blot analysis of Tet2 and Fbp1 levels in livers from the HFD mice infected with AAV8 or AAV8-*shTet2* for 10 days, and treated with or without metformin (300 mg/kg/day) for another 10 days as indicated. (**F**, **G)** GTT (F) and ITT (G) were performed in the HFD mice infected with AAV8 or AAV8-*shTet2* for 10 days, and treated with or without metformin (300 mg/kg/day) for another 10 days as indicated (n = 6).

## Discussion

Our study revealed the function of TET2 in the regulation of gluconeogenesis. Notably, gluconeogenesis levels in *TET2* knockout HepG2 cells and primary mouse hepatocytes significantly decreased even under glucagon treatment, demonstrating that TET2 is essential for glucagon-induced upregulation of glucose output. Our findings link the novel function of TET2 to the gluconeogenic process. However, the precise role of TET2 in the pathophysiology of type 2 diabetes (T2D) remains unclear and requires further research. Additionally, the mechanisms by which TET2 is upregulated during gluconeogenesis deserve further exploration.

A recent study showed that metformin treatment can reduce hepatic glucose production by targeting FBP1 (3), suggesting that FBP1 may be a promising target for diabetes management. However, the regulation of FBP1 remains poorly understood, despite several studies indicated that the promoter methylation mediates FBP1 expression silencing in cancer (20, 21). Here, we found that TET2 positively regulates FBP1 in response to glucagon treatment, indicating that targeting TET2 may represent a promising strategy for treating T2D.

We further investigated how TET2 specifically regulates FBP1 expression in response to glucagon stimulation. Like most chromatin-modifying enzymes, which require DNA sequence-specific binding proteins, such as DNA transcription factors, for recruitment and regulation of specific gene expression (22), TET2 also requires binding partners to modulate particular pathways in a context-dependent manner. This is supported by findings that Wilms tumor protein (WT1) (23) and Smad Nuclear Interacting Protein 1 (SNIP1) (24) are indispensable for TET2 to suppress leukemia cell proliferation and regulate the cellular DNA damage response, respectively. Our data demonstrated that TET2 upregulates FBP1, thereby enhancing gluconeogenesis in an HNF4α-dependent manner. Furthermore, metformin-induced phosphorylation of HNF4α at Ser 313 impairs its binding ability to TET2 and reduces TET2 recruitment to the FBP1 promoter, leading to decreased FBP1 expression. Importantly, TET2-FBP1 inhibition has a synergistic effect with metformin in HFD mice.

Intriguingly, a clinical investigation study revealed that TET2 mutations occur more frequently in the diabetes mellitus (DM) group than the non-DM, suggesting a potential connection between TET2 and insulin resistance (25). Additionally, inactivating mutations of epigenetic regulator TET2 led to metabolic dysfunction, including clonal hematopoiesis, and aggravate age- and obesity-related insulin resistance in mice (26). Regarding HNF4α variants, an analysis utilizing exome sequencing data demonstrated that human genetic variations in HNF4α disrupted its protein structure and function, impaired insulin secretion, reduced sensitivity to insulin, and increased the risk of T2D in individuals (27, 28). Furthermore, it has been reported that a point mutation in FBP1 can reduce the efficacy of metformin treatment (3), providing a genetic evidence for the role of TET2-FBP1 axis in the therapeutic effect of metformin. In summary, our findings uncovered a previously unknown function of TET2 in gluconeogenesis. TET2, together with HNF4α, facilitates FBP1 expression by maintaining the hypomethylation of FBP1 promoter, which leads to increased gluconeogenesis. Thus, targeting HNF4α-TET2-FBP1 axis may represent a promising strategy to lower blood glucose in T2D.

## Materials and Methods

### Cell Culture and Transfection

HepG2, HEK293T, and primary mouse hepatocytes were cultured in DMEM medium supplemented with 10% fetal bovine serum (FBS) and incubated at 37°C in a 5% CO_2_ atmosphere. For plasmids or siRNA transfection, FuGENE HD (#E2311, Roche, WI, USA) and Lipofectamine 2000 (#11668030, Invitrogen, CA, USA) were utilized respectively. For lentivirus production, polyethyleneimine (PEI) from EZ Trans (#AC04L091, Heyuan Liji, Shanghai, China) was used. All transfection procedures were conducted according to the manufacturers’ instructions. The sequences of HNF4α siRNA are listed as follows: 5’-AAUGUAGUCAUUGCCUAGGTT-3’ and 5’-UCUUGUCUUUGUCCACCACTT-3’.

### Isolation of Primary Mouse Hepatocytes

Primary mouse hepatocytes were isolated by perfusing the liver with pre-warmed Hank’s Balanced Salt Solution (HBSS) at a flow rate of 5-7 mL/minute for 5 minutes after anesthetizing the mouse. Once the liver appeared pale, the perfusion solution was switched to a digestion buffer, maintaining the same flow rate and duration. The liver was then transferred to a 10 cm dish containing 10 mL of digestion medium. Hepatocytes were released by gently cutting and agitating the liver, after which 20 mL of cold PBS was added to halt the trypsin digestion. The cell suspension was filtered through a 70 μm nylon mesh (BD Falcon), centrifuged at 50 g for 2 minutes, and the cell pellet was washed twice with cold PBS at 50g for 5 minutes each. The hepatocytes were then ready for subsequent experiments.

### Western Blot and Immunoprecipitation

Cell lysates were prepared using NP40 lysis buffer, proteins were separated by SDS-PAGE. The following primary antibodies were used: Tubulin (#66031-1-Ig, Proteintech, Wuhan, China; 1:1000), TET2 (#18950S, CST, MA, USA; 1:1000), HNF4α (#32591, SAB, MD, USA; 1:1000), HNF4α-pS313 (ab78356, Abcam, Cambridge, UK, 1:1000), FBP1 (#HPA005857, Sigma, MO, USA; 1:1000), and HA (#3724, CST, MA, USA; 1:1000). Secondary antibodies included anti-rabbit (#L3012, SAB, MD, USA; 1:1000) and anti-mouse (#L3032, SAB, MD, USA; 1:1000). For immunoprecipitation, total protein was incubated with protein A beads (#37484, CST, MA, USA) and TET2 antibody (#MABE 462, Millipore, MO, USA; 1:1000) for 3 hours at 4°C, followed by western blot analysis.

### Reverse transcription qPCR

Total RNA was extracted using an RNA purification kit (#B0004D, EZBioscience, MN, USA). Real-time PCR was conducted using SYBR Green (#11200ES03, YEASEN, Shanghai, China) following cDNA synthesis. β-actin was used as an internal control. The sequences of the qPCR primers are as follows:

hTET2-forward: GATAGAACCAACCATGTTGAGGG;

hTET2-reverse: TGGAGCTTTGTAGCCAGAGGT;

hFBP1-forward: CGCGCACCTCTATGGCATT;

hFBP1-reverse: TTCTTCTGACACGAGAACACAC;

hACTIN-forward: CATGTACGTTGCTATCCAGGC;

hACTIN-reverse: CTCCTTAATGTCACGCACGAT;

mTet2-forward: AGAGAAGACAATCGAGAAGTCGG;

mTet2-reverse: CCTTCCGTACTCCCAAACTCAT;

mFbp1-forward: CACCGCGATCAAAGCCATCT;

mFbp1-reverse: CCAGTCACATTGGTTGAGCCA;

mActin-forward: GTGACGTTGACATCCGTAAAGA;

mActin-reverse: GCCGGACTCATCGTACTCC.

### Chromatin immunoprecipitation (ChIP)

ChIP assay was performed based on a manufacturer’s instructions (#P2078, Beyotime Biotechnology, Shanghai, China). Briefly, HepG2 cells were washed with PBS and cross-linked using formaldehyde of final concentration of 1% (Sigma, USA) for 10 min, at room temperature. Genomic DNA of HepG2 cell lysate was digested with micrococcal nuclease (#D7201S, Beyotime Biotechnology, Shanghai, China) for 20 min at 37 ℃, and then sonicated to produce nucleotide fractions of 200–1000 base pairs. The mixed solution was centrifuged at 13,000 rpm at 4°C to separate the sediments. The sediments were resuspended with ChIP dilution buffer (#P2078, ChIP Assay Kit, Beyotime Biotechnology, Shanghai, China) and incubated with anti-TET2 antibody (#18950S, CST, MA, USA; 1:1000), anti-5mC antibody (#HA601350, HUABIO, Hangzhou, China), anti-5hmC antibody (#HA601351, HUABIO, Hangzhou, China), and anti-IgG primary antibody [#10284-1-AP, Proteintech, Wuhan, China] at 4°C overnight. The mixture was incubated with Protein A/G Sepharose beads (#HY-K0230, MCE, Shanghai, China) for 2 hours at 4°C, followed by centrifugation at 1000 rpm at 4 °C. The antibody-conjugated Protein A/G Sepharose beads were washed with PBS, and then eluted with fresh elution buffer, to dissociate the conjugated DNA. DNA was isolated using a DNA purification kit (#B518141, Sangon Biotech, Shanghai, China), followed by ChIP-PCR. The primers used for amplification of immunoprecipitated DNA are as follows:

FBP1-Forward 5ʹ-GATCCCCGACCTTGTCTGAA-3ʹ;

FBP1-Reverse 5ʹ-TCGCGGAAACCTTTAGACGC-3ʹ.

### Gene Knockout Cells and Mutagenesis

The CRISPR/Cas9 system was employed to generate TET2 knockout cells. Lentiviruses were produced by co-transfecting HEK293T cells with plasmids containing sgRNA (8μg), psPAX2 (6μg), and pMD2.G (2μg) in a 10 cm dish. The collected supernatant was used to infect the target cells. Following selection with puromycin, Western blot analysis was performed to assess the efficiency of the knockout. The targeting sequence for TET2 sgRNA was 5’-GATTCCGCTTGGTGAAAACG-3’. For mutagenesis, human HNF4α cDNA was cloned into a pLVX-2×Flag lentiviral expression vector. Both of site-directed mutants of HNF4α, including HNF4α-S313A and HNF4α-S313D, were generated by PCR using KOD FX (#KFX-201, TOYOBO, Japan) following the manufacturer’s protocol. Briefly, HNF4α plasmids were amplified, and the products were digested with DpnI enzyme (#R0176V, New England Biolabs, MA, USA) before being transformed into NcmDH5-α (#MD101-1, NCM Biotech, Suzhou, China) for amplification.

### Immunofluorescence

For immunofluorescence, the HepG2 cells or mouse liver sections were incubated with related primary antibodies overnight at 4°C. Mouse anti-TET2 (#ANM4475, IPODIX, Wuhan, China; 1:100) and rabbit anti-HNF4α (#ET1611-43, HUABIO, Hangzhou, China; 1:100) were used. The cells or sections were rinsed with phosphate-buffered saline (PBS) for 3 times, then incubated with Alexa 488-conjugated or Alexa 555-conjugated secondary antibodies (#A0460, # A0423, Beyotime, Shanghai, China; 1:200) for 2 h at room temperature. The HepG2 cells or sections were rinsed with PBS for 3 times and stained with ProLong™ Diamond Antifade Mountant with DAPI solution (#P36971, Invitrogen, CA, USA), and coated with coverslips (#174950, Invitrogen, CA, USA). All images were captured by using a confocal microscope and processed with Fluoview software (Olympus-FV3000, Tokyo, Japan), ImageJ and Photoshop CS5.

### Glucose Production

Cells were washed three times with PBS before the medium was replaced with glucose-free DMEM (#A14430-01, Gibco, CA, USA) supplemented with 20 mM sodium lactate and 2 mM sodium pyruvate. After an 8-hour incubation, glucose levels in the supernatants were measured using a glucose assay kit (#A22189, ThermoFisher, MA, USA).

### PTT, GTT and ITT

For the PTT and GTT, mice were fasted for 16 h and 12 h, respectively, before receiving an intraperitoneal (i.p.) injection of either pyruvate (2g/kg) or glucose (2g/kg). In the ITT, mice fasted for 4 h in the morning prior to receiving an insulin injection (1U/kg, i.p.). Blood glucose levels were measured at intervals of 0, 15, 30, 45, 60, 90, and 120 minutes after injection using a glucometer during tail vein bleeding.

### Animal

All animal experiments were approved by the Animal Care and Use Committee at Fudan University (Shanghai, China). Male C57BL/6J mice, aged 6 weeks, were obtained from the Charles River Labs. Homozygotes of the whole-body *Tet2* knockout (Tet2 *KO*) strain were originally purchased from the Jackson Laboratory (Jackson Laboratories, Bar Harbor, ME, stock No. #:023359). These mice were housed under specific-pathogen-free (SPF) conditions at 25°C with a 12-hour light/dark cycle and were fed either a standard chow or a high-fat diet (HFD) (Research Diets, 45% calories from fat). To induce insulin resistance, wild-type C57BL/6J mice aged 6 weeks were placed on an HFD for 11 days or 12 weeks. For the liver-specific *Tet2*-knockdown mice, we transduced AAV8 shRNA against *Tet2* to knock down *Tet2* in the liver (designed and synthesized by Hanbio, Shanghai, China) of mice. Scramble shRNA (AAV8) was used as a negative control. Target sequences for *Tet2* shRNA was 5’-GGAAUAUCCCACAUGAAAGGCAGCC-3’. In brief, male C57BL/6J mice aged 6 weeks were switched from standard chow to HFD diet. Mice were then randomly divided into four groups: AAV8, AAV8 + metformin, AAV8-*shTet2*, AAV8-*shTet2* + metformin. AAV8 and AAV8-*shTet2* were diluted with saline to 1×10^12^ vector genomes/ml, and 100 μL was injected through the tail vein for each mouse.

### Statistical Analysis

Sample numbers are indicated in the figures and figure legends. The Student’s *t* test was used to determine the significant difference between two groups, and one-way analysis of variance (ANOVA) with Tukey’s post hoc test for multiple comparisons when more than two groups were analyzed. Data analysis was performed using SPSS version 16.0 (SPSSInc., Chicago, IL, USA) or GraphPad Prism (version 8.0.2.; GraphPad Software, Inc., San Diego, CA, USA). A p-value of less than 0.05 was considered statistically significant. *, **, *** and **** indicated statistically significant results compared to the control and represented P < 0.05, P < 0.01, P < 0.001, and P < 0.0001, respectively.

## Acknowledgments

This work was supported by the National Key R&D Program of China (2022YFA0807100, 2020YFA0803400/2020YFA0803402), the National Natural Science Foundation of China (82372754, 82472850, 82172936, 81972620, 82121004 and 82073128), the Shanghai Rising-Star Program (24QA2709900), the Program for Professor of Special Appointment (Eastern Scholar) at Shanghai Institutions of Higher Learning and the Fundamental Research Funds for the Central Universities.

